# A computational examination of the two-streams hypothesis: which pathway needs a longer memory?

**DOI:** 10.1101/2020.09.30.321299

**Authors:** Abolfazl Alipour, John Beggs, Joshua Brown, Thomas W. James

## Abstract

The two visual streams hypothesis is a robust example of neural functional specialization that has inspired countless studies over the past four decades. According to one prominent version of the theory, the fundamental goal of the dorsal visual pathway is the transformation of retinal information for visually-guided motor behavior. To that end, the dorsal stream processes input using absolute (or veridical) metrics only when the movement is initiated, necessitating very little, or no, memory. Conversely, because the ventral visual pathway does not involve motor behavior (its output does not influence the real world), the ventral stream processes input using relative (or illusory) metrics and can accumulate or integrate sensory evidence over long time constants, which provides a substantial capacity for memory. In this study, we tested these relations between functional specialization, processing metrics, and memory by training identical recurrent neural networks to perform either a viewpoint-invariant object classification task or an orientation/size determination task. The former task relies on relative metrics, benefits from accumulating sensory evidence, and is usually attributed to the ventral stream. The latter task relies on absolute metrics, can be computed accurately in the moment, and is usually attributed to the dorsal stream. To quantify the amount of memory required for each task, we chose two types of neural network models. Using a long-short-term memory (LSTM) recurrent network, we found that viewpoint-invariant object categorization (object task) required a longer memory than orientation/size determination (orientation task). Additionally, to dissect this memory effect, we considered factors that contributed to longer memory in object tasks. First, we used two different sets of objects, one with self-occlusion of features and one without. Second, we defined object classes either strictly by visual feature similarity or (more liberally) by semantic label. The models required greater memory when features were self-occluded and when object classes were defined by visual feature similarity, showing that self-occlusion and visual similarity among object task samples are contributing to having a long memory. The same set of tasks modeled using modified leaky-integrator echo state recurrent networks (LiESN), however, did not replicate the results, except under some conditions. This may be because LiESNs cannot perform fine-grained memory adjustments due to their network-wide memory coefficient and fixed recurrent weights. In sum, the LSTM simulations suggest that longer memory is advantageous for performing viewpoint-invariant object classification (a putative ventral stream function) because it allows for interpolation of features across viewpoints. The results further suggest that orientation/size determination (a putative dorsal stream function) does not benefit from longer memory. These findings are consistent with the two visual streams theory of functional specialization.

## 1 Introduction

One of the oldest ideas in neuroscience is functional specificity. According to this idea, each neuron or group of neurons (i.e., cortical column, brain region, etc.) primarily subserves one function. This is contrasted with another equally old idea in neuroscience, distributed processing, where neurons or brain regions engage in a variety of different functions. Moreover, functional specificity maintains that it is possible for multiple brain regions to be the primary contributors to a specific function and, also, distributed processing maintains that one brain region can be equally important for multiple functions. For example, in the view of functional specialization, multiple brain regions can be the primary contributors to language comprehension, but a region that primarily contributes to language comprehension cannot also be a primary contributor to mathematical abilities (Fedorenko et al., 2011). It is widely accepted that functional specialization exists in early sensory regions, where receptive field properties such as retinotopic or tonotopic maps provide clear examples. However, even a relatively low-level visual area like V4, has been shown to have a variety of functions, rather than a primary function (Desimone & Schein, 1987), despite early attempts to classify V4 as more functionally specific and only a color processor (Zeki, 1980). Thus, the presence of functional specialization in higher-level visual areas or even in more cognitive brain regions is clearly a topic of debate (Connolly et al., 2012; Haxby et al., 2000, 2014; Kanwisher, 2000, 2010; Mahon & Cantlon, 2011; Zerilli, 2017)

As with most dichotomies in science, it is likely that functional specialization is better represented by a spectrum — from highly specialized to highly distributed — with the functional profiles of different brain regions falling somewhere along the spectrum, rather than simply landing at either end. Theorizing a spectrum, however, raises further questions, fundamental among them being what characteristics of functions require (or allow) them to be more (or less) specialized or distributed across brain regions? One candidate characteristic is the incompatibility of the computations performed – if functions are computationally compatible, then multiple functions could presumably share the architecture of a given brain region, whereas a function that is not computationally compatible with others may require specialized circuitry. An example of this is the proposed sharing of the architecture in the visual lateral occipital cortex (area LO) of humans for object recognition by vision and touch (Amedi et al., 2001, 2002; James et al., 2002). Sensory systems are usually considered specialized in neuroscience, but there is considerable evidence that area LO shares circuitry between two (if not three) sensory systems (Amedi et al., 2007; James et al., 2011; Lacey et al., 2009; Pascual-Leone & Hamilton, 2001) suggesting that this putatively specialized visual brain region distributes its processing across different sensory systems. This overlap is deemed possible in LO because object recognition by vision and touch share similar computations of volumetric shape.

An excellent example of functional specialization that falls in this debated part of the spectrum is the influential two visual streams hypothesis/theory (TVSH). According to this hypothesis, visual information processing splits into two cortical streams after its initial processing in early visual areas. In colloquial terms, the function of the ventral visual stream is to shape our visual perception (what) and, according to Milner and Goodale’s (1991, 2008) version of the TVSH, the function of the dorsal stream is to act on the environment or visually guide our movements (how) (Milner & Goodale, 2008).

Although what differentiates Milner and Goodale’s version of the TVSH from others is that the defining characteristic of each stream is its output modality – either perception or action – the TVSH also ascribes other processing characteristics to each stream that serve that specific output modality. One of those characteristics is memory. If the fundamental goal of the dorsal visual pathway is the transformation of retinal information for visually-guided motor behavior, then it should process inputs using absolute (or veridical) metrics and the processing should occur at the moment when the movement is initiated, necessitating very little, or no, memory. Conversely, because the ventral visual pathway does not involve motor behavior (interaction with the veridical world), the ventral stream should process inputs using relative (or illusory) metrics and accumulate or interpolate sensory evidence over long time constants or, in other words, have a long memory.

The purpose of this study was to investigate the extent to which the dorsal and ventral streams may be functionally separated due to the incompatibility of their computations. As cited above, the dorsal and ventral streams differ computationally on many dimensions, but here we focused on the combination of differences in metrics and memory. We tackled this goal using computational modeling and, specifically, by using recurrent neural network models that had explicit memory parameters. We trained the models on two tasks. A *viewpoint-invariant object classification* task was used to assess ventral stream processes, because solving it relies on relative metrics and benefits from the accumulation of sensory evidence. An *orientation/size determination* task was used to assess dorsal stream processes, because solving it relies on absolute metrics and can be accomplished accurately in the moment. We hypothesized that neural networks trained on these two tasks would use different amounts of memory, suggesting that separation of these processes in the brain may be necessary for optimal performance. Additionally, we hypothesized that the longer memory in ventral stream is driven by temporal interpolation of object features, and we tested this in two ways. First, when objects are rotated about their vertical axis, they occlude their features (self-occlusion). Therefore, viewpoint-invariant object classification will need to have a longer memory to interpolate features from multiple viewpoints when self-occlusion happens. Therefore, without self-occlusion, the need for long memory should diminish. Secondly, the reliance on feature interpolation should increase when the features of objects in a class are more visually similar and should decrease if the objects in a class are not chosen based on visual similarity. In other words, if visually dissimilar, but semantically linked objects are placed in one class, having a longer memory will not help the viewpoint-invariant object classification as much as if objects in a class are visually similar.

## 2 Methods

### 2.1 Datasets

#### 2.1.1 COIL-100 Dataset

The first dataset that we used for evaluating the memory span differences between object and orientation tasks was the Columbia Object Image Library (COIL-100) dataset (Nene et al., 1996). This dataset contains images of 100 natural objects. In each object class, there are 72 images of the same object from different viewpoints in 5-degree increments rotated about the vertical axis (i.e., yaw).

In the COIL-100 dataset, we derived an orientation task according to the following procedure: first, 46 objects in the COIL-100 dataset that show ambiguous orientations upon rotation were removed (bottles, cans, cups), and the remaining 54 object classes were used. Subsequently, the following six orientation classes were defined:

- Class 1: All orientations of 0°±5° (5-degree leeway) and their mirror orientations (180°±5°) regardless of the object identity.
- Class 2: All orientations of 30°±5° (5-degree leeway) and their mirror orientations (210°±5°) regardless of the object identity.
- Class 3: All orientations of 60°±5° (5-degree leeway) and their mirror orientations (240°±5°) regardless of the object identity.
- Class 4: All orientations of 90°±5° (5-degree leeway) and their mirror orientations (270°±5°) regardless of the object identity.
- Class 5: All orientations of 120°±5° (5-degree leeway) and their mirror orientations (300°±5°) regardless of the object identity.
- Class 6: All orientations of 150°±5° (5-degree leeway) and their mirror orientations (330°±5°) regardless of the object identity.

Each of the six classes had 324 images (a total of 1,944 images). Note that changes in object orientation in this dataset always accompanied changes in the apparent 2-D size of objects as well, and for this reason, we considered the “orientation task” to be an orientation/size determination task.

For consistency across tasks, the object task used the same 54 objects that were used in the orientation task. In each of the 54 object classes, there were 36 images of the same object from different viewpoints. These viewpoints were images of the objects with 30°±5° degree increments. The 30° increment was used to be consistent with the previous literature on the limits of viewpoint invariant object recognition in humans (Bülthoff et al., 1995; Bülthoff & Edelman, 1992). This resulted in an object task with 1,944 images, matching the total number of images in the orientation task.

**Fig. 1.**
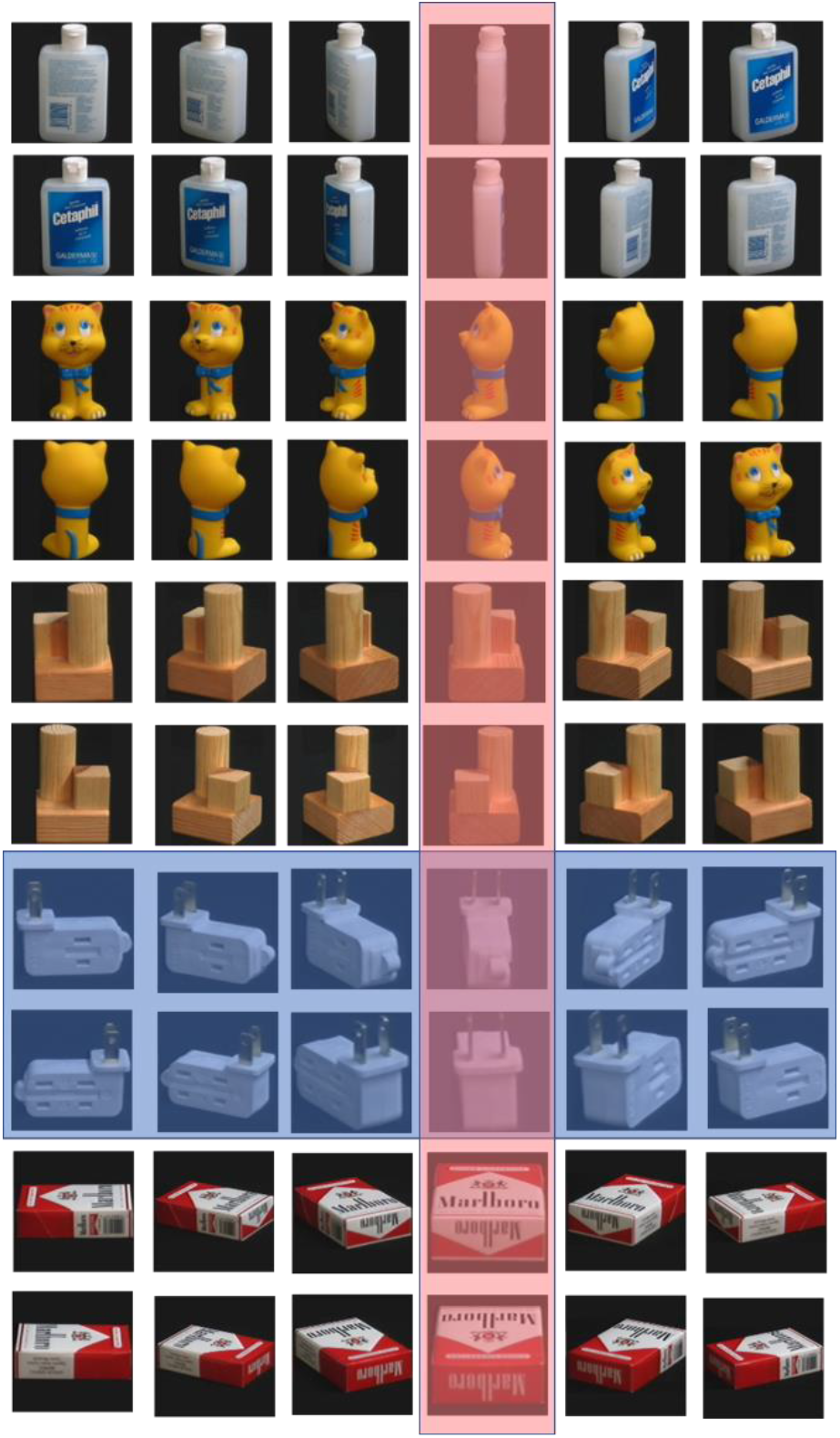
Example images from the COIL-100 dataset. Shown are a subset of 5 objects (of 54 total) at 6 orientations and their mirror images (12 of 36 total images in a class). Columns represent 30 degrees of rotation. Rows are the five different objects, with one of a pair of rows showing the six unique orientations and the other of the pair showing the mirror images. The training dataset also included orientations at ±5° to either side of the orientations shown (out of 36 possible orientations). The blue shaded images represent a single object class, and the red shaded images represent a single orientation class (10 of 324 total images in a class).

#### 2.1.2 MIRO dataset

A different dataset was needed to investigate the effect of rotating about the visual axis (i.e., pitch) and also to investigate the effect of visual similarity within an object class. We chose the Multi-view Images of Rotated Objects (MIRO) dataset introduced by (Kanezaki et al., 2018). The MIRO dataset contains several object categories and images of the different instances of each category at many orientations, created by rotating about the visual axis.

The relatively large number of instances per category in the MIRO dataset compared to the COIL-100 dataset allowed us to create two pairs of object/orientation tasks. For the “visual” task pair, the object task had 22 object classes with high visual similarity within each class, and the orientation task had 8 classes. More specifically, for the visual object task, images were placed within object classes based on both visual and semantic similarity. Therefore, images in each class belonged to the same semantic class (e.g., all were images of buses), and they had similar visual features as well (e.g., only buses of similar type and size –such as school buses– were placed in one class). This resulted in 22 object classes in the visual object task. Note that with rotations about the visual axis, the size of the 2-D image does not significantly change when orientation is changed. Therefore, in contrast to the orientation task with the COIL-100 dataset, the orientation task items from MIRO dataset were driven purely by changes in orientation rather than changes in both orientation and size. More importantly, rotation along the visual axis did not create self-occlusion of objects, and all features were present in every frame. This allowed us to test if the need to retain features from previous frames was driving memory span in our networks. The visual orientation task was to classify the orientation of images regardless of their object identity. In the orientation task, the eight classes were as follows:

- Class 1: All orientations of 0° regardless of the object identity.
- Class 2: All orientations of 45° regardless of the object identity.
- Class 3: All orientations of 90° regardless of the object identity.
- Class 4: All orientations of 135° regardless of the object identity.
- Class 5: All orientations of 180° regardless of the object identity.
- Class 6: All orientations of 225° regardless of the object identity.
- Class 7: All orientations of 270° regardless of the object identity.
- Class 8: All orientations of 315° regardless of the object identity.

For the “semantic” task pair, the object task had 6 object classes with low visual similarity within each class and the orientation task had 4 classes. More specifically, for the semantic object task, images were placed within object classes solely based on their semantic similarity (they did not need to be visually similar). Therefore, images in each class belonged to the same semantic class (e.g., all were images of buses), but they had varied visual features (e.g., all buses, regardless of their type, size, etc. were placed in one class). The semantic orientation task was to classify the orientation of images regardless of their object identity. Number of classes were reduced in the semantic orientation task to ensure similar difficulty between the object and orientation tasks for the semantic task pair. Using the semantic pair of tasks allowed us to see if featural similarity within an object class was another factor that drives memory span. In other words, if feature interpolation contributes to the need for longer memory spans, reducing within-class featural similarity should reduce memory dependence. For the semantic orientation task, the four orientation classes were as follows:

- Class 1: All orientations of 0° and their mirror orientations (180°) regardless of the object identity.
- Class 2: All orientations of 45° and their mirror orientations (225°) regardless of the object identity.
- Class 3: All orientations of 90° and their mirror orientations (270°) regardless of the object identity.
- Class 4: All orientations of 135° and their mirror orientations (315°) regardless of the object identity.

**Fig. 2.**
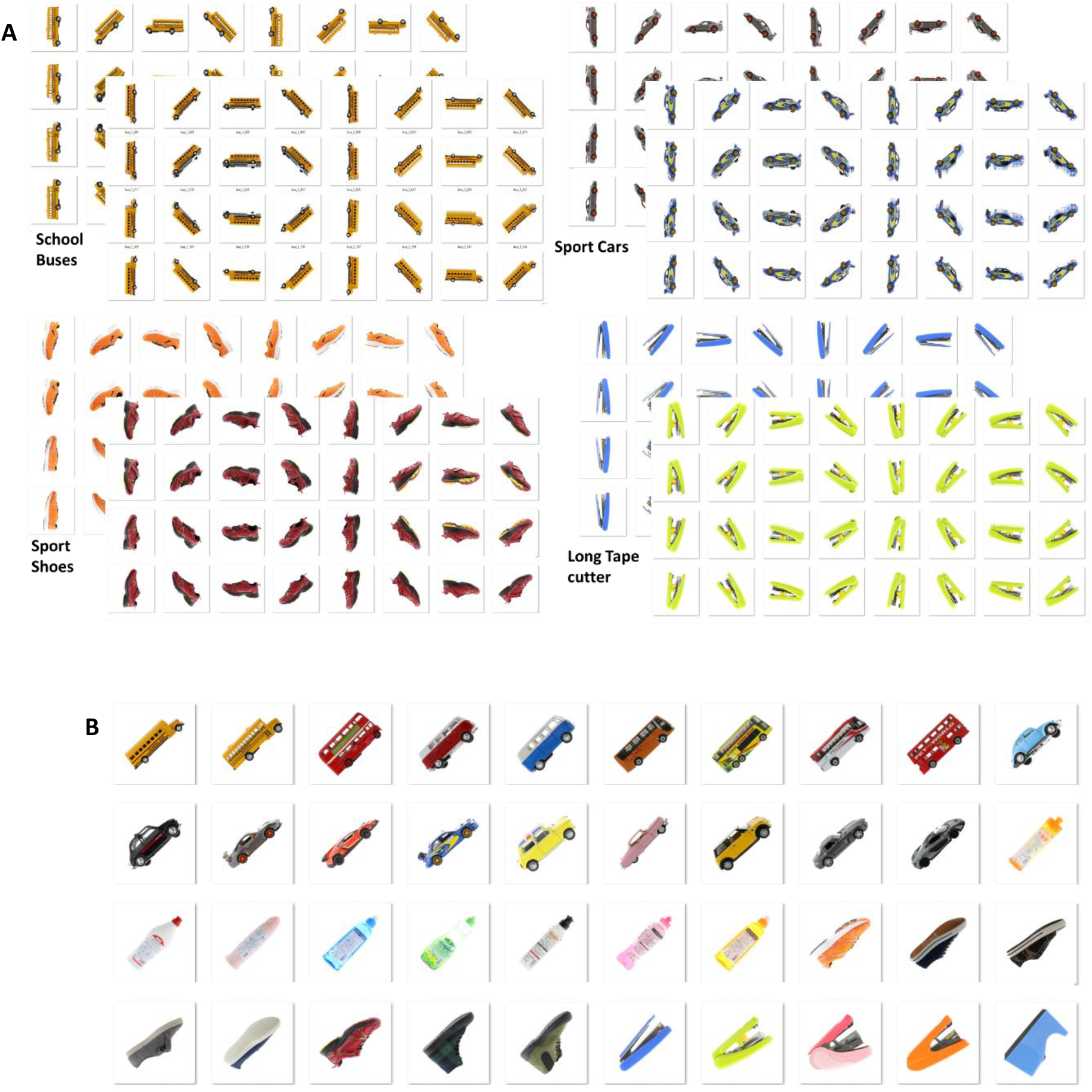
Example images from the visual task pair derived from the MIRO dataset. A: Four object classes from the semantic object task (of the 22 total classes). Shown are all 64 images that exist in each object class. In this pair of tasks, instances within an object class were chosen based on visual AND semantic similarity. B: example of 40 images within the 45° orientation class (out of 114). Other images that are not shown were rotated around their *x* axis for 16 degrees (i.e., rolled 16 degrees). An example of such rotations on an image is shown in Figure 3.C. Note that mirror images were considered separate classes in the visual MIRO pair of tasks.

**Fig. 3.**
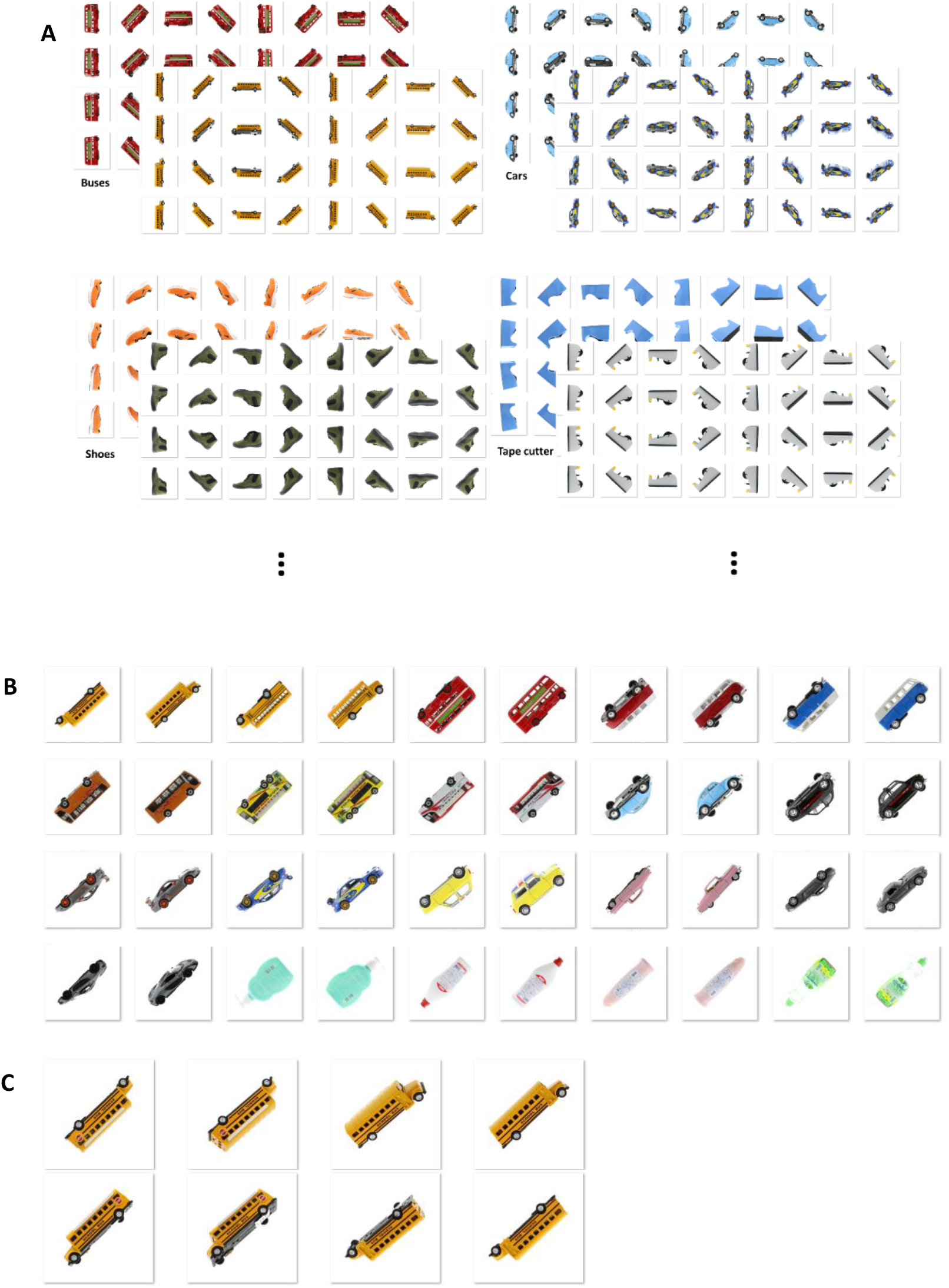
Example images from the semantic task pair derived from the MIRO dataset. A: Four object classes from the semantic object task (of the six total). Shown are 64 images out of 320 total images in each object class. Other images that are not shown were rotated around their *x* axis for 16 degrees (i.e., rolled 16 degrees). In this pair of tasks, instances within an object class were chosen based on semantic class, regardless of visual similarity. B: 40 example images from the 45° and 225° orientation class (of 480 total) used in the semantic orientation task. C: Examples of rolled objects. The viewpoints shown in the first 4 images of B are rolled either 16° or −16°.

**Fig. 4.**
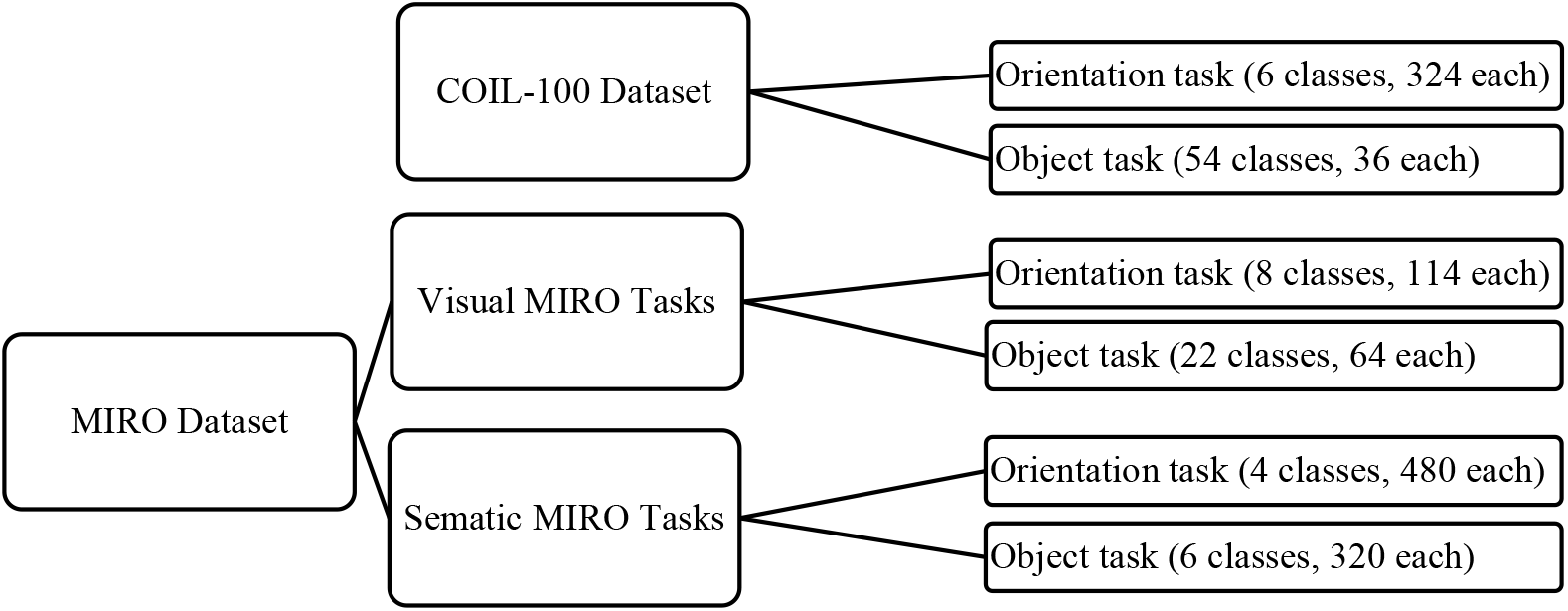
Organization of the datasets and tasks.

### 2.2 Networks and training procedure

Because convolutional neural networks (CNNs) have shown a reasonable similarity to the primate visual system (Cadieu et al., 2014; Schrimpf et al., 2018; D. L. Yamins et al., 2013; D. L. K. Yamins & DiCarlo, 2016), we used them as approximations of the early visual areas in both streams. This is in line with recent computational studies on the visual system that have shown the ability of CNNs to provide interesting insights into the functions of the visual system (Bao et al., 2020; Cadieu et al., 2014; Kar et al., 2019; Lotter et al., 2020).

To this end, we started with two architecturally identical CNNs (details in figure 6) and trained one of them on the viewpoint-invariant object classification task and the other on the orientation/size determination task. Multiple instances of feedforward networks were trained on each of the two tasks. From the trained feedforward networks, we selected one for each task by matching their test accuracies. The weights in these trained feedforward networks were then frozen, and the frozen-weight networks were used in the next training/testing phase. This was done based on previous studies showing the effectiveness of transfer learning (Tan et al., 2018).

#### 2.2.1 Using RNNs and sequences of images to measure memory dependence in each task

Next, to see if temporal information (i.e., memory) in a sequence of images was important for the network’s performance, we replaced the last layer of the trained, frozen-weight, early networks with a recurrent neural network (RNN), either a long-short term memory (LSTM) network (experiment 1) or a leaky-integrator echo state network (LiESN, experiment 2), as shown in figure 6. Because the RNN part of each model could generate outputs based on both current and past input images, we were able to train the models on sequences of images and measure the effect of retaining information from the past images (memory) in each task. Hence, the task of the RNNs was to *classify images within a sequence of images*. For the object task, the stimulus was a sequence of images that contained randomly drawn viewpoints of the same object in each training/test epoch. For the orientation task, the stimulus was a sequence of images that contained randomly drawn objects with the same orientation in each training/test epoch. In a training epoch, the model was presented a sequence and the task was to classify each image using the previous and the current image. Experiments were performed using PyTorch (*Automatic Differentiation in PyTorch | OpenReview*, n.d.) running on python 3.6.

### 2.3 Experiment 1: using LSTMs

In experiment 1, we replaced the last layer of each CNN with an LSTM and trained one model on the object task and the other on the orientation task. The LSTM network was implemented based on the original LSTM paper (Hochreiter & Schmidhuber, 1997).

With the weights in the early CNN frozen, the two ways for the network to improve accuracy on the tasks during learning were adjustment of the memory coefficient or adjustment of the output weights. The influence of adjustments of the output weights on accuracy is related to network size. Because the purpose of the study was to examine the role of memory, we wanted to limit the interaction between changes in memory coefficient and changes in output weights. In other words, we wanted to limit the possibility that larger RNNs were producing memory effects (due to intrinsic recurrence) even with low memory coefficients. Therefore, the first phase of each experiment was to find the network size(s) where the network size had minimal impact on the relationship between memory coefficient and accuracy.

Therefore, in experiment 1, each model was a combination of a trained, frozen-weight CNN with an LSTM RNN replacing its last layer. Each model was further trained (i.e., the RNN part of the model was trained) on three pairs of object/orientation tasks, one pair from the COIL-100 dataset and two pairs from the MIRO dataset. The training was done using sequences of randomly drawn images, as described above. The training duration for each task was 200 epochs. During the 200 epochs, changes in memory coefficient were recorded as accuracy reached its plateau. Alternatively, when a given network reached an accuracy of more than 99% before 200 epochs, we stopped the training and recorded the memory coefficient at 99% accuracy. To obtain statistical reliability, this simulation was repeated 30 times, and the average memory coefficients along with their corresponding test accuracies were recorded for every 10 epochs. This procedure enabled us to observe the optimization of memory coefficient as models were being trained on object/orientation tasks from the three pairs of tasks.

To measure how much of the past information (history, memory) is relevant to the classification performance in an LSTM network, we used the ratio of the forget gate to the input gate as an indicator of network reliance on the past information. The forget gate value in an LSTM network is a number between 0 and 1, and it is multiplied by the hidden state value to control the effect of the past information (hidden states) on the current output of the LSTM network. The input gate value in an LSTM network is also a number between 0 and 1, and it is multiplied by the current input of the LSTM network to indicate how much the current inputs should affect the cell state of an LSTM network. Accordingly, the ratio of the forget gate to the input gate can show how much a given network relies on its memory (hidden states) compared to its current inputs to generate an output. The larger this ratio, the more reliance there is on the past information (memory) in the LSTM network. We call this ratio the ‘LSTM memory coefficient’. To obtain this memory coefficient, we trained our LSTM network on each of the tasks. Subsequently, we fed the test data to the network and measured the logarithm of the ratio of *forget gate values to input gate values* in the LSTM network as images were passing through the network. The average logarithm of (forget gate/input gate) for the entire test dataset was calculated and reported for both object and orientation tasks. This procedure was used for all three task pairs.

### 2.4 Experiment 2: using LiESNs

In experiment 2, we used LiESN RNNs instead of LSTM RNNs to test if the results were independent of the type of RNN used.

LiESNs are a variant of Echo state networks (ESNs). ESNs are recurrent networks with randomly connected units that have sparse connectivity. The input weights (weights from the previous layer) and the recurrent weights in ESNs are randomly initialized and kept fixed afterward. The output nodes are supposed to adjust their weights to derive the desired computations from the “reservoir of computations” that the ESN network provides. This architecture was first introduced by Jaeger (Jaeger, 2004, 2007) as a brilliant but straightforward alternative to remedy the problems that come with backpropagation in RNNs. The simplicity of ESNs, their relative biological plausibility, and their capability to approximate highly nonlinear functions make them useful models for examining the role of recurrence in memory function.

LiESNs are ESNs with leaky units, which essentially allows the ESN to produce a running average with the history set by a parameter. In a typical LiESN, the dynamics of the network with N_U_ inputs and N_R_ reservoir units are governed according to:

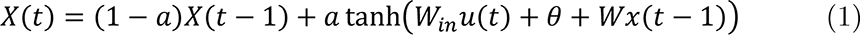

Where *X* is the state of the reservoir with N_R_ dimensions, *t* is the time, *u* is the input signal with N_U_ dimensions, *Win* is the input weight matrix (N_U_ by N_R_), *Wx* is the reservoir weight matrix (N_R_ by N_R_), and *θ* is the bias term. The parameter *a* is the leaking rate (Gallicchio et al., 2017; Jaeger, 2001; Schaetti et al., 2016). Leaking rate varies from 0 to 1 and, as Equation 1 shows, when the leaking rate is zero, the state of the reservoir depends solely on its past state and the current inputs are ignored, but when the leaking rate is 1, the state of the reservoir depends solely on the current inputs and its past state is ignored.

The purpose of the current study was to examine the influence of memory on the object and orientation tasks. To do that, we modified the typical LiESN by decoupling the influence of leaking rate on the current inputs and on the past state. Instead, the influence of the current inputs was kept constant, while a “memory” parameter was used to vary the history used in the running average. This changed produced the following equation:

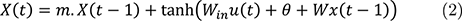

Equation 2 makes explicit that the value of the current inputs is no longer modified by a leaky rate parameter, and the value of the past state is modified by a “memory coefficient”, m. The value of m has a different effect on the state of the reservoir than the leaky rate. When m is 1, the state of the reservoir is determined by a balance of the current inputs and past inputs, but when m is 0, the state of the reservoir is determined solely by the current inputs. Thus, larger values of *m* indicate longer memory.

In experiment 2, the training procedure was the same as experiment 1 with the difference that training was done only for 18 epochs. Additionally, each of the 30 simulations were initialized by a randomly chosen memory coefficient between 0 and 1.

**Fig. 5.**
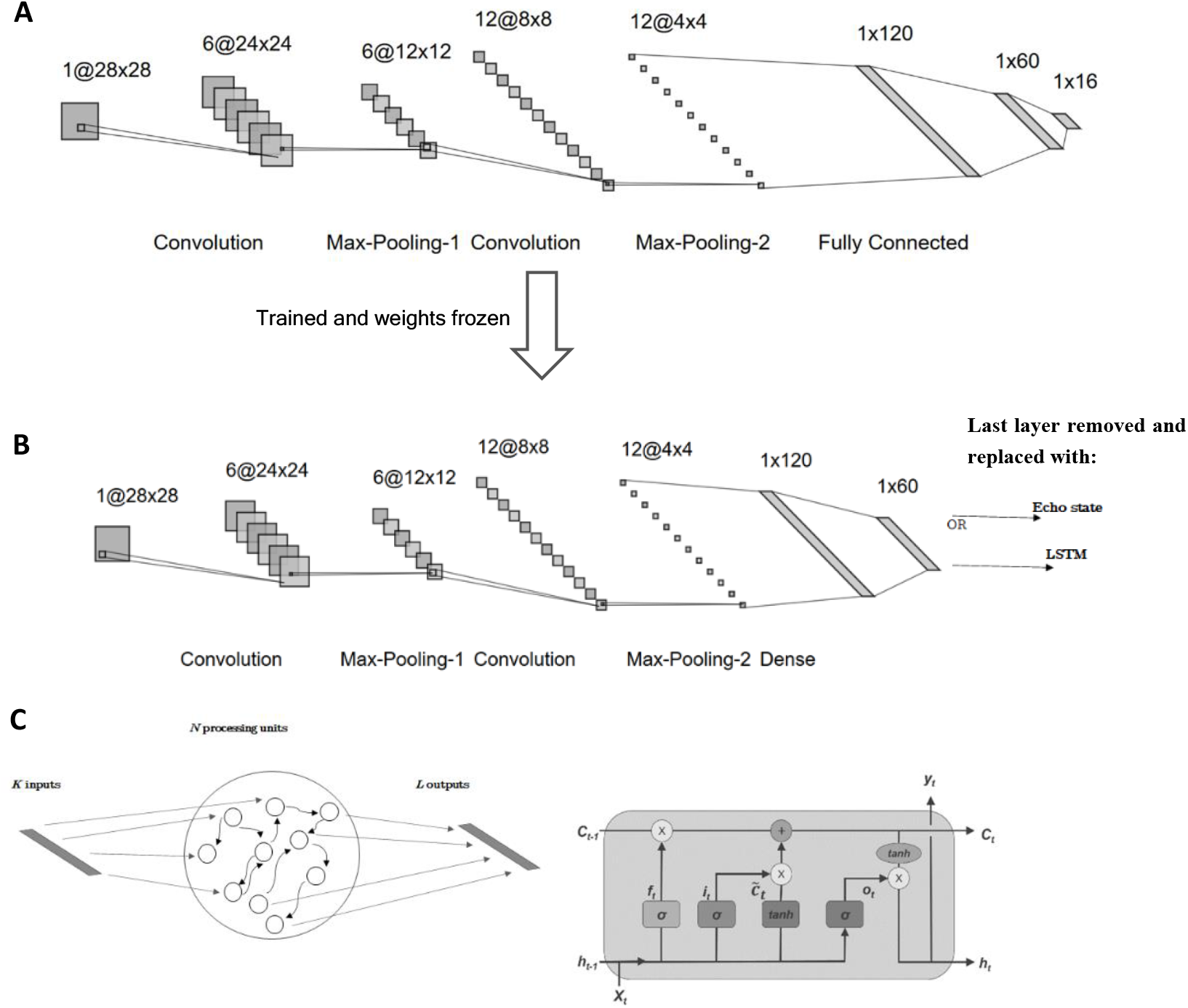
The architecture of the model. A: Early CNN architecture. The early levels of the network comprised two convolutional layers with max-pooling and three fully connected layers. Note that fully connected layers were not recurrent but were connected to all nodes in the previous layer. B: The last layer in the classifier was replaced with either an LSTM RNN (Exp 1) or an echo state RNN (Exp 2). C. Left: Schematic representation of an echo state network. Right: schematic representation of an LSTM network (from (Zheng et al., 2017), under Creative Commons Attribution License).

## 3 Results

### 3.1 Feedforward convolutional networks (without recurrent networks)

We trained many feedforward CNNs (figure 3.) to either perform viewpoint-invariant object classification (object task) or orientation/size determination (orientation task) on the three pairs of tasks. After 50 epochs of training with a learning rate of 10^-4^, the performance of the feedforward CNNs was tested using the test dataset. For each pair of task, we selected a pair of CNNs that produced similar accuracies, one with the object task and one with the orientation task. It is worthwhile noting that the accuracy for the orientation task was higher across all of the simulations, which is why we selected pairs of CNNs for the next stage that produced similar accuracy on the two tasks. Because the weights on the chosen CNNs were frozen in the subsequent training, the input to the RNNs was derived from CNNs that produced similar accuracies. See table 1 for details of the accuracies in each pair of tasks.

**Table 1.** Numerical values of classification accuracies in feedforward networks.

|  | COIL-100 Dataset |  | MIRO Semantic |  | MIRO Visual |  |
| --- | --- | --- | --- | --- | --- | --- |
|  | Object | Orientation | Object | Orientation | Object | Orientation |
| Accuracy | 61% | 59% | 70% | 71% | 47% | 51% |

### 3.2 Experiment 1: LSTM

We observed that overall, LSTM network size has a negative correlation with memory coefficient and a positive correlation with accuracy for both tasks, suggesting the presence of a memory-network size tradeoff (see supplementary figures 1-3). We assume this happened because larger LSTMs have more parameters/weights to adjust to solve the tasks. Because the model could solve the tasks by adjusting either the memory coefficient (reliance on previous states) or the model weights, we needed to identify network sizes that made the tasks appropriately challenging and where there was a high reliance on memory for both tasks. Therefore, for comparison, we chose the smallest network size (n=20) to compare memory across tasks because memory reliance was the highest in small networks.

As described in the Methods, in the second stage of training, the CNN part of each model was weight-frozen while they were trained with sequences of images from the three pairs of tasks shown in Figure 5. The RNN part of each model learned the sequences by adjusting its weights through gradient descent. As mentioned in the methods section, the LSTM memory coefficient was defined as the logarithm of the forget gate to input gate ratio. We found that the LSTM memory coefficient was higher in the object task in the COIL-100 dataset where objects are rotated about their vertical axis and self-occlusion happens (*M* = 0.83, *SEM* = 0.066 for object task compared to *M* = 0.30, *SEM* = 0.014 for orientation task, *t*(59) = −7.67, *p* = 1.45e-10). Additionally, memory coefficient for the object task was still higher in the Visual MIRO pair of tasks (Figure 9.B) where objects had visually similar features but were rotated about their visual axis which resulted in no self-occlusion (*M* = 0.60, *SEM* = 0.036 for object task compared to *M* = 0.173, *SEM* = 0.017 for orientation task, *t*(59) = −10.67, *p* = 2.66e-15). We noted that the gap between memory coefficient between object and orientation tasks was slightly lower in the Visual MIRO pair of tasks compared to COIL-100, supporting our hypothesis that self-occlusion contributed to longer memory coefficient in the object task.

There was no significant difference between the object and orientation tasks in the Semantic MIRO task pairs where objects were rotated around their visual axis and did not share many similar visual features (Figure 9.C) (*M* = 0.26, *SEM* = 0.017 for object task compared to *M* = 0.20, *SEM* = 0.022 for orientation task, *t*(59) = −1.97, *p* = .054). We also looked at the value of memory coefficient for the object task in Visual MIRO and compared it to the object task in the Semantic MIRO (Fig 9.D) and found that there was a significant difference between the two conditions (*M* = 0.60, *SEM* = 0.036 Visual MIRO object compared to *M* = 0.26, *SEM* = 0.017 for Semantic MIRO object task, *t*(59) = −8.63, *p* = 5.50e-12). This suggested that feature similarity was driving the need for longer memory in the object task. Moreover, this effect was not sensitive to the number of classes (supplementary figure 1). This observation supported our hypothesis that: first, the object task requires longer memory, and second, self-occlusion and featural similarity partially drive memory span.

**Fig. 6.**
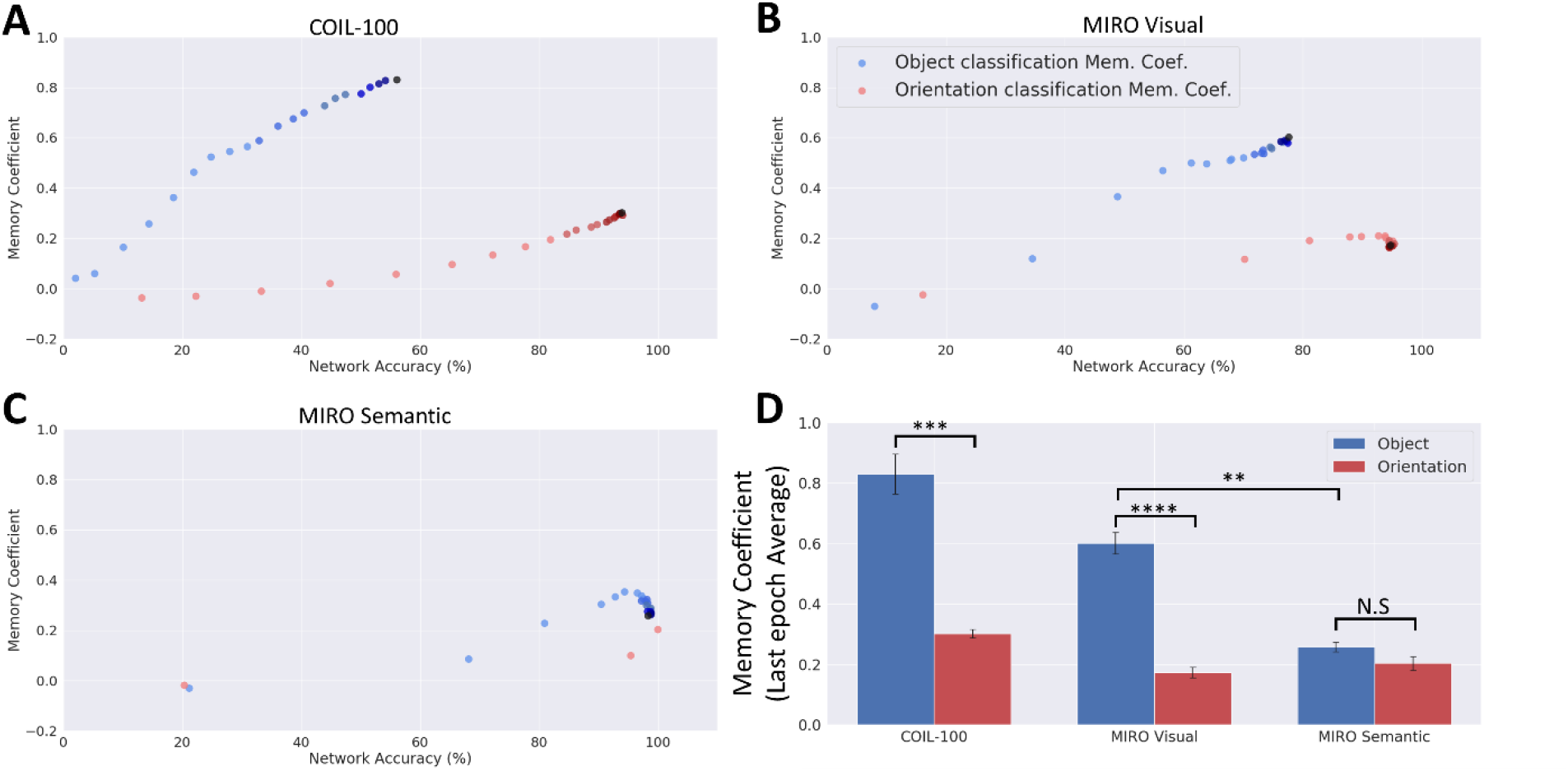
Memory coefficient as a function of task accuracy and task type across learning epochs. Dots represent every tenth epoch (of 200), with darker shades indicating later epochs. A) Results from the COIL-100 dataset. B) Results from the MIRO Visual dataset. C) Results from the MIRO Semantic dataset. D) Memory coefficient on the last epoch as a function of dataset and task type. Error bars reflect the standard error of the mean (SEM). Note that we stopped the training in MIRO semantic after epoch 30 since the network reached 99% test accuracy.

### 3.3 Experiment 2: LiESN

Similar to LSTMs, we observed that there was a negative correlation between network size and accuracy in LiESNs. We assume this happened because larger LiESNs have a more diverse repertoire of computations and, more importantly, there are more output weights (i.e., more parameters) to adjust to solve the tasks. Therefore, like with LSTMs, it was imperative that we identified network sizes that made the tasks appropriately challenging and where there was at least some reliance on memory for both tasks. Accordingly, we first searched to find network sizes for which the output weight adjustment did not differentially affect the adjustment of the memory coefficient in either of the tasks. To that end, we analyzed the similarity of the accuracy-memory coefficient relationships across tasks in multiple network sizes.

In contrast to LSTMs, we found that small LiESNs did not necessarily provide the best testbeds for comparing the two tasks. In two out of the three task pairs, there were dissimilarities between the memory-accuracy profiles in the two tasks when networks were too small or too large (supplementary figures 5-7). Instead, the memory-accuracy profile was most similar across tasks for the COIL-100 dataset when the network size was 160, for the MIRO Semantic dataset when the network size was 20, and for the MIRO Visual dataset when the network size was 160 (see supplementary figure 5-7). These network sizes were used for all further simulations in experiment 2.

As described in the Methods, in the second stage of training, the CNN part of each model was weight-frozen while they were trained with sequences of images from the three pairs of tasks shown in Figure 4. The RNN part of each model learned the sequences by adjusting both the memory coefficient and the output weights by gradient descent.

Figure 7 shows changes in accuracy and memory coefficient across learning epochs for both tasks. For the COIL-100 dataset, accuracy and memory coefficient increased across epochs for both tasks, and as hypothesized, the memory coefficient was adjusted higher for the object task than for the orientation task. (*M* = 0.76, *SEM* = 0.005 for object task compared to *M* = 0.52, *SEM* = 0.044 for orientation task, *t*(59) = 5.32, *p* = 1.72e-6)

**Figure 7.**
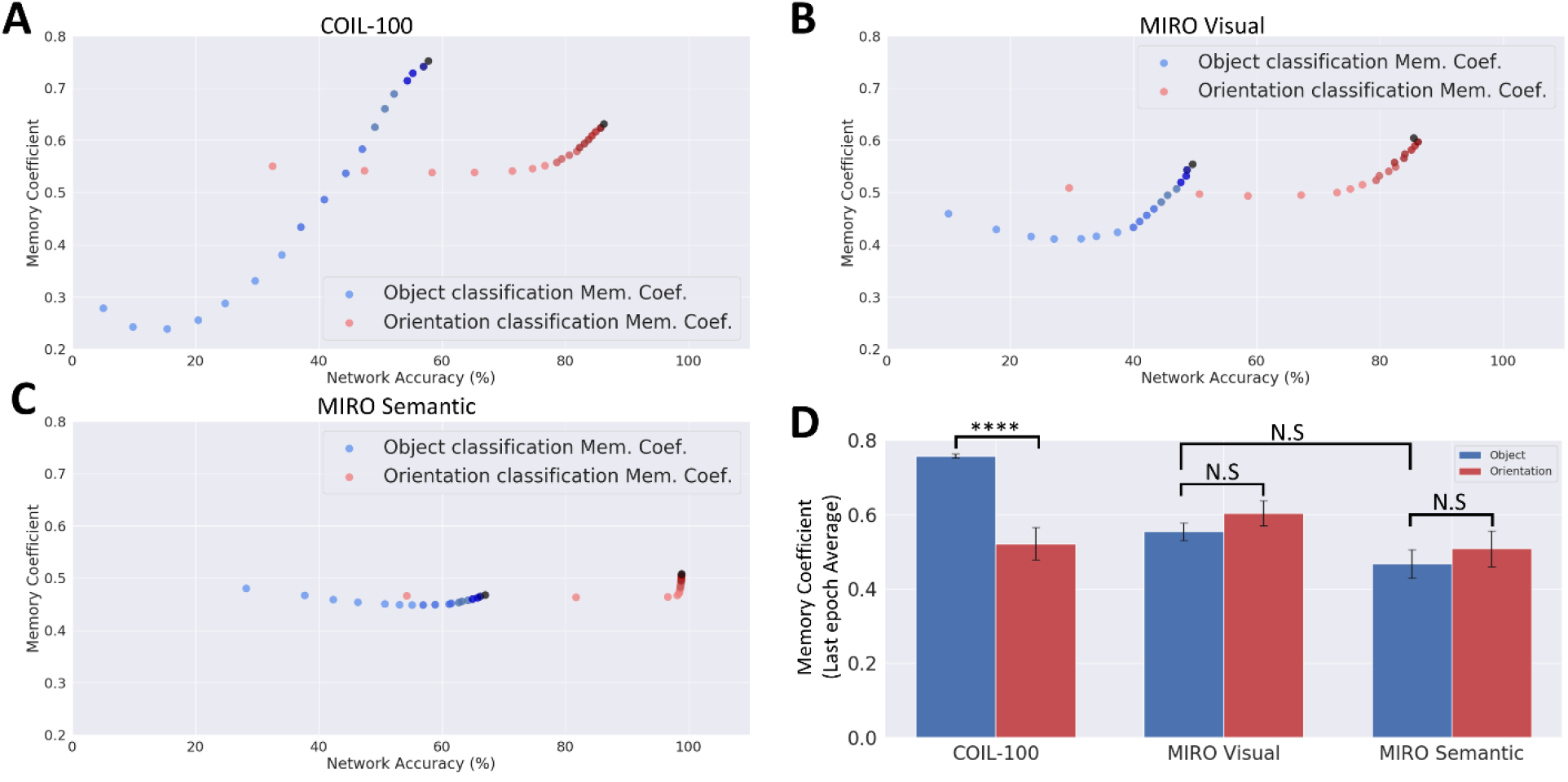
Memory coefficient as a function of task accuracy and task type across learning epochs. Dots represent every epoch (of 18), with darker shades indicating later epochs. A) Results from the COIL-100 dataset. B) Results from the MIRO Visual pair of tasks. C) Results from the MIRO Semantic pair of tasks. D) Memory coefficient on the last epoch as a function of task pair and task type. Error bars reflect the standard error of the mean (SEM).

For the MIRO datasets, accuracy again increased across epochs. However, changes in memory coefficient were different from the COIL-100 dataset. The Visual and Semantic Orientation tasks required approximately the same amount of memory as COIL-100, but the Visual and Semantic Object tasks did not see the same large increase in memory. That is, for the MIRO dataset, there was no significant difference between Object and Orientation tasks for either the Visual or Semantic datasets. (MIRO visual: *M* = 0.55, *SEM* = 0.024 for object task compared to *M* = 0.60, *SEM* = 0.034 for orientation task, *t*(59) = −1.22, *p* = .23) (MIRO Semantic: *M* = 0.47, *SEM* = 0.038 for object task compared to *M* = 0.51, *SEM* = 0.048 for orientation task, *t*(59) = −0.65, *p* = .52) This difference between COIL-100 and MIRO datasets may have been due to the hypothesized difference in processing required by rotations about the vertical axis (COIL-100) versus rotations about the visual axis (MIRO). However, that would not explain the differences between results with LiESN and LSTM model results.

Another difference between LiESNs and LSTMs was their sensitivity to the number of classes in the object and orientation tasks. Because of the way the object and orientation tasks were developed from the datasets, the “standard” object tasks always had more classes than the “standard” orientation tasks. To ensure that the greater memory for the object task was not simply due to the object task having more classes, we ran simulations where the number of classes in the object tasks was lowered to be the same as the number in the corresponding orientation tasks. LSTM models were insensitive to this manipulation, producing similar results regardless of class size (see supplementary figure 4). However, LiESNs were highly sensitive to class size (see Figure 8). First, in LiESN models, the memory effect between object and orientation tasks with the COIL-100 dataset with different class sizes (see Fig 8.A) vanished when the class sizes were equated (*M* = 0.48, *SEM* = 0.034 for object task compared to *M* = 0.51, *SEM* = 0.039 for orientation task, *t*(59) = −0.70, *p* = .49) and, second, a memory effect that did not exist with the Visual MIRO Visual task pair with different class sizes (see Fig 8.B) appeared when the class sizes were equated. (*M* = 0.47, *SEM* = 0.035 for object task compared to *M* = 0.63, *SEM* = 0.037 for orientation task, *t*(59) = −3.16, *p* = .0025). There was no change with the Semantic MIRO task pair.

**Figure 8.**
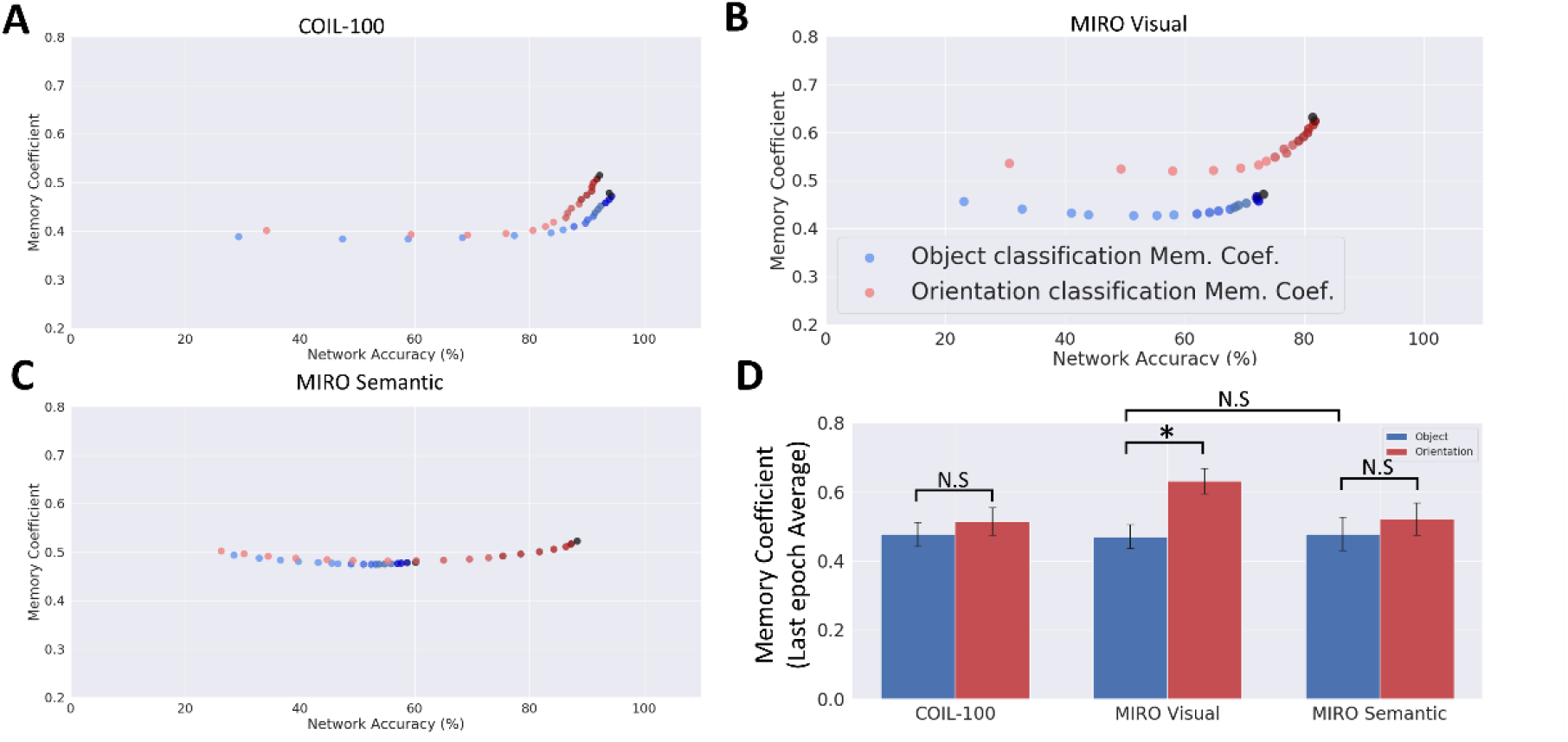
Memory coefficient as a function of task accuracy and task type across learning epochs when the number of classes across task types were equated. Dots represent every epoch (of 18), with darker shades indicating later epochs. A) Results from the COIL-100 dataset. B) Results from the MIRO Visual dataset. C) Results from the MIRO Semantic dataset. D) Memory coefficient on the last epoch as a function of the dataset and task type. Error bars reflect the standard error of the mean (SEM).

## 4 Discussion

Consistent with the TVSH, we showed that model visual systems ending in LSTM RNNs rely more heavily on memory when performing a viewpoint-invariant object classification (object task) compared to an orientation/size determination (orientation task). According to the TVSH, the object task taps functions that are considered part of the ventral visual stream, and the ventral visual stream is also considered to computationally rely more on memory than the dorsal stream. To investigate the factors that give rise to longer memory in object tasks, we analyzed two additional factors. First, as hypothesized, the self-occlusion of objects that were rotated about their vertical axis (in depth) relied more heavily on memory. Our rationale for this finding is that ‘remembering’ visual features from previous viewpoints in a random sequence is more helpful when those features change across viewpoints, as they do when self-occlusion occurs. Second, as hypothesized, restricting object classes to only visually similar images, as opposed to semantically similar (but possibly visually dissimilar) images, also relied more heavily on memory. Our rationale for this finding is that ‘remembering’ visual features from previous viewpoints in a random sequence is more helpful when features in the class are visually similar because remembering features is only useful if it helps to classify ambiguous images in the future. Overall, our results on LSTM networks suggest that reliance on memory is beneficial for the object task by allowing the accumulation or interpolation of object features. The orientation task did not show this same benefit, which is consistent with another hypothesis of the article, that the dorsal and ventral streams are anatomically isolated because their distinct functions are computationally incompatible.

Although model visual systems ending in LSTM RNNs were highly consistent with the stated hypotheses, models ending in LiESN RNNs were not as consistent. First, in LiESN models, changing the number of classes changed the memory dependence for both tasks. Our rationale for this effect is that, in LiESNs, the number of output classes determines the number of plastic weights, and each additional class adds N additional weights to the model (where N is the number of hidden units in the LiESN). According to our size-accuracy analysis (Supplementary figure 5-7), increasing network parameters had a complicated relationship with the accuracy and memory in LiESNs. Second, the LSTM memory coefficient was defined as an average of individual forget to input gate ratios, which we consider a more fine-grained measure of memory than the memory coefficient in LiESNs, which was a general property of the entire network. Therefore, the major difference between the LSTM and LiESN models to mimic the TVSH pattern may have been that individual weights were plastic in LSTMs, whereas recurrent weights were frozen in LiESNs.

One may think the length of image sequences could drive the differences in memory lengths in these tasks in each class of the dataset, i.e., object task networks might have longer memory simply because they have been learning longer sequences of images. However, we deliberately used longer sequences in the orientation classes of all three datasets. Notably, there were 324 images in each orientation class compared to 36 in each object class in the COIL-100 dataset. Similarly, we had 64 and 180 images in object and orientation classes for the Visual MIRO dataset, respectively. For the Semantic MIRO dataset, there were 320 images in each object class and 480 images in each orientation class.

Alternatively, one may think that the total number of images in each task may drive the memory dependence in our networks. In other words, the reliance on memory may be reflecting the amount of information that is being learned in each task. However, in COIL-100 and MIRO pair of tasks, the number of images in each task were equal (1944, 1440, and 1920 in COIL-100, Semantic MIRO, and Visual MIRO, respectively).

One interesting effect that we observed was the inverse relationship between network size and memory dependence. Both LiESN and LSTM models showed an inverse relationship between network size and their need for larger memory spans. Moreover, we observed that memory dependence decayed differently for each task, and orientation classification can be less sensitive to network size on some occasions (e.g., supplementary figure 7). Therefore, the number of parameters in a network (network size), dataset size, and memory coefficient can contribute to network performance, and their effect should be considered before drawing conclusions. While the relationship between network architecture and memory capacity in echo state networks has been explored elsewhere (Gallicchio et al., 2018), the exact relationship between network size and memory capacity deserves further investigation.

From another perspective, our results were in line with clinical findings in patients and the two-streams hypothesis for primate vision. Meanwhile, It is noteworthy to mention that some studies called initial assumptions of the two-streams hypothesis into question (Hesse & Schenk, 2014; Konen & Kastner, 2008; Rogers et al., 2009), (see (Schenk & Hesse, 2018) for a critical review). Thus, our results cannot generalize to the entire dorsal pathway. In particular, it is essential to note that according to Kravitz et al., the dorsal stream itself comprises three different sub-pathways: the parieto-prefrontal, the parieto-medial temporal, and the parieto-premotor paths (Kravitz et al., 2011). Specifically, the parieto-prefrontal path is involved in spatial working memory and eye movement control, the parieto-medial temporal pathway is critical for spatial navigation and spatial long-term memory, and the parieto-premotor pathway is heavily involved in visually controlled movements such as reaching and grasping (Kravitz et al., 2011). Due to the nature of the tasks we used here, our results are most relevant to the parieto-premotor pathway.

The very short-term nature of memory in the dorsal pathway has been extensively studied using psychophysical and imaging studies and generated mixed results. For example, Cant et al. showed that while naming objects can be primed, a grasping movement cannot be primed by previous grasping movements (Cant et al., 2005), supporting the short-term nature of the visuomotor control representations. Similarly, Jax and Rosenbaum reported that while priming in a visually guided obstacle avoidance task with short delays between the priming stimulus and actual task was possible, this effect went away with delays that were longer than a second (Jax & Rosenbaum, 2007). Meanwhile, the idea that in delayed motor-controlled tasks, the source of information switches from dorsal to ventral pathway was challenged by some studies (Schenk & Hesse, 2018). For example, in a study by Himmlbach et al., a bilateral optic ataxia patient (IG) showed strong activity in regions around his lesion in the dorsal pathway in both immediate and delayed reaching tasks (Himmelbach et al., 2009). Moreover, the famous patient D.F. showed that she could perform delayed visually guided movement as well as controls when the environmental cues (allocentric information) were *not* available (Hesse & Schenk, 2014). This study lent credit to the idea that the dorsal pathway can still keep information related to visually guided behavior for long delays (>2seconds), and it is the contextual information that becomes available after a delay. Moreover, Singhal et al. used functional magnetic resonance imaging (fMRI) and showed that during delayed action tasks, the dorsal stream retains rough plans for actions, and right before the movement, it extracts fine-grained information from the ventral stream (Singhal et al., 2013). Adding to the complexity of the matter, van Elk et al. showed that immediate *grasping* movement engages ventral stream areas to a larger extent, as compared to *pointing* movement as evident by electroencephalography (EEG) signals (van Elk et al., 2010). Therefore, the movement control system for both immediate and delayed actions recruit regions in the ventral streams as well, and the delineation between dorsal-ventral streams is not very strict.

The very short memory span of the dorsal stream could be understood from the perspective of its inputs as well. Magnocellular inputs mostly innervate the dorsal pathway, whereas the ventral pathway is more innervated by parvocellular inputs (Merigan & Maunsell, 1993). The main difference between these two types of inputs is that the magnocellular cells are better at the classification of higher temporal frequencies, whereas parvocellular cells are more suitable for higher spatial frequencies. Additionally, magnocellular cells are 20 milliseconds faster regarding their response latency to stimuli (Bullier & Nowak, 1995). The faster dynamics of the dorsal pathway is in line with what we found in our study.

Because a similar two-stream dissociation is suggested in other sensory modalities such as somatosensation (Dijkerman & de Haan, 2007; James & Kim, 2010) or audition (Hickok & Poeppel, 2007; Rauschecker, 2018), the relationship between short term memory and motor control tasks might even go beyond vision. However, our current datasets and tasks are limited to vision, and specific tasks for each modality would be required to find out if such a relationship holds for other sensory modalities as well.

Another issue is the fact that the majority of the computational models of the visual cortex are based on the ventral stream (e.g., (Lotter et al., 2016, 2020)), while models incorporating both ventral and dorsal pathways are less common (but see (O’Reilly et al., 2017, 2020) as an example incorporating both streams). We believe that newer models of the visual cortex should incorporate the tasks relevant to both streams. This will demonstrate if different tasks (e.g., object recognition vs. motor control) require incompatible computational paradigms that need different circuitries.

From a broader perspective, our results suggest that some computations may require separate circuitry to serve their purpose, as we hypothesized in the introduction. Accordingly, this *computational incompatibility* might be one of the reasons for functional specificity in the brain. In particular, our results bear some similarities with studies on lateralization of spatial frequency processing in the brain. In spatial information processing, high spatial frequencies are processed more efficiently in the left hemisphere, while low spatial frequencies are processed more efficiently in the right hemisphere (Flevaris & Robertson, 2016; Howard & Reggia, 2007; Jager & Postma, 2003). Here, it seems that memories are retained on longer time scales in the ventral stream while their retention is much shorter in the dorsal stream, a counterpart for the lateralization of information processing (Figure 15). This is in line with previous reports on circuit timescales in other species (Murray et al., 2014; Siegle et al., 2021)

**Fig. 15.**
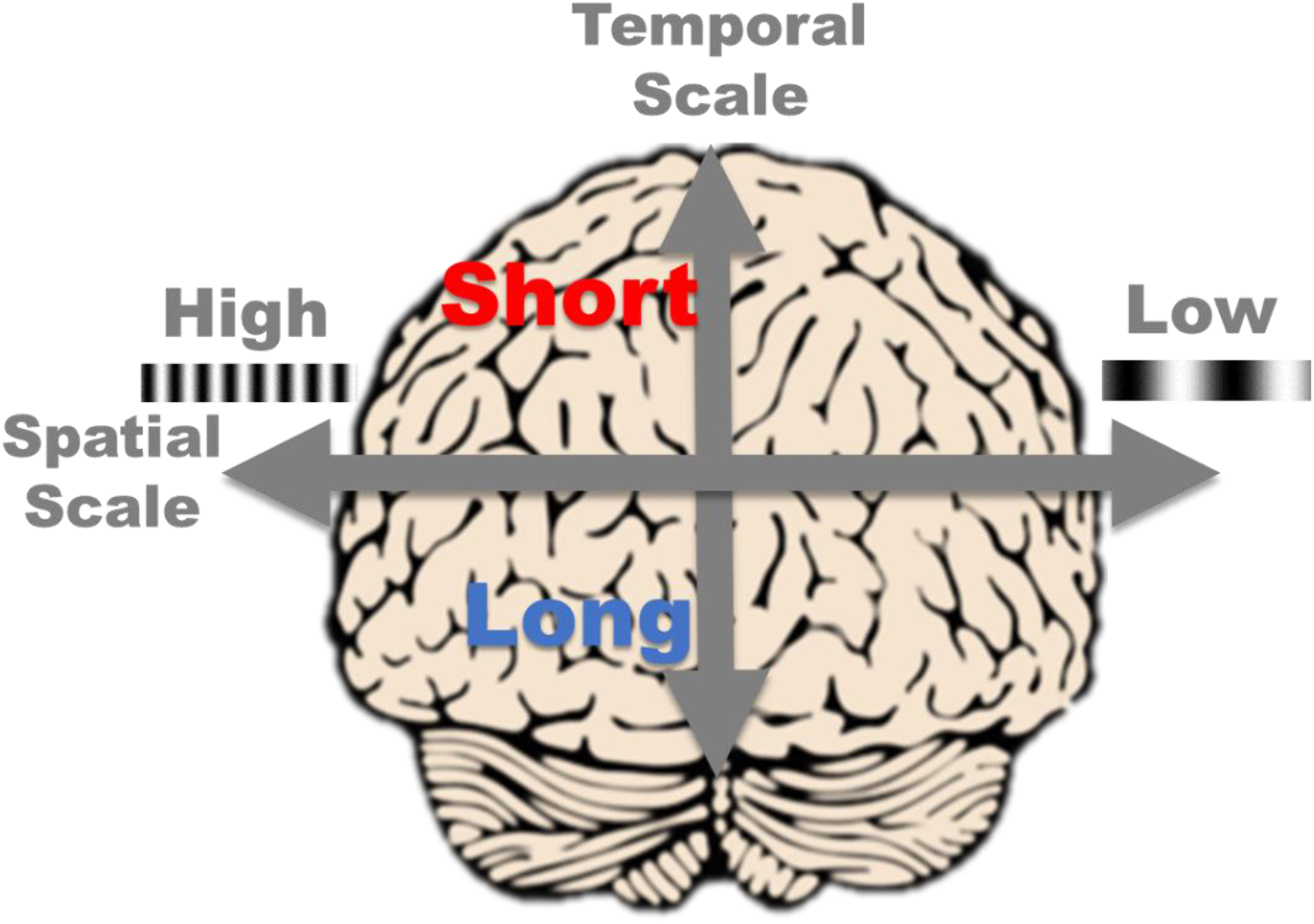
Localization of temporal and spatial information processing in the brain. One example is the lateralization of spatial frequency processing across the two hemispheres. Another example is the dorsal-ventral stream differences in their memory span: dorsal stream keeps information on shorter time scales compared to ventral stream. Our results suggest that the dorsal-ventral memory span differences might have been caused by the computational incompatibility of the tasks they perform.

## 5 Limitations

Our current study comes with several shortcomings. First, we could have added a motor control task to our orientation classification task to be able to make a direct comparison between our model and the parietal-premotor pathway. However, using such a task would make it challenging to create architecturally identical networks and compare their memory coefficients.

Second, the dorsal pathway receives information from subcortical sources such as superior colliculus and these sources are mostly innervated by magnocellular cells with higher sensitivity to lower spatial frequencies. We could have used larger-sized kernels for convolutional layers that we used to approximate the dorsal pathway. However, doing so would render the comparison of recurrent networks less straightforward.

Finally, even though orientation classification is a prerequisite for visually guided movements, in principle, it can happen in the ventral stream as well. We believe that such orientation classification in the ventral stream emerges as a byproduct of object recognition, while it is one of the core computations in the dorsal stream. Therefore, training networks to exclusively perform orientation classification might have ameliorated this issue, but it is not a perfect model of the dorsal stream.

## 6 Conclusions

In the present study, we showed that object classification needs a longer memory compared to orientation classification in LSTMs. We used multiple RNN architectures and datasets to investigate this effect, and our results suggest that this memory dependence is driven by the usefulness of accumulating visual features when self-occlusion and featural similarity are included in the viewpoint-invariant object classification task. Future studies should use sophisticated motor control tasks to study this hypothesis. Furthermore, it is possible to study this effect in other modalities in which a two-stream dissociation has been proposed (i.e., somatosensation and hearing).

## 7 Acknowledgment

This research was supported in part by Lilly Endowment, Inc., through its support for the Indiana University Pervasive Technology Institute and in part by the Indiana METACyt Initiative. The Indiana METACyt Initiative at IU was also supported in part by Lilly Endowment, Inc. The authors acknowledge the Indiana University Pervasive Technology Institute for providing Carbonate HPC resources that have contributed to the research results reported within this paper (Stewart et al., 2017).

## 8 Supplementary Material

### 8.1 Code

The code used in this study is available at (https://github.com/Abolfazl-Alipour/DorsalAndVentralMemoryDifferences).

### 8.2 Supplementary figures

**Supplementary figure 1:**
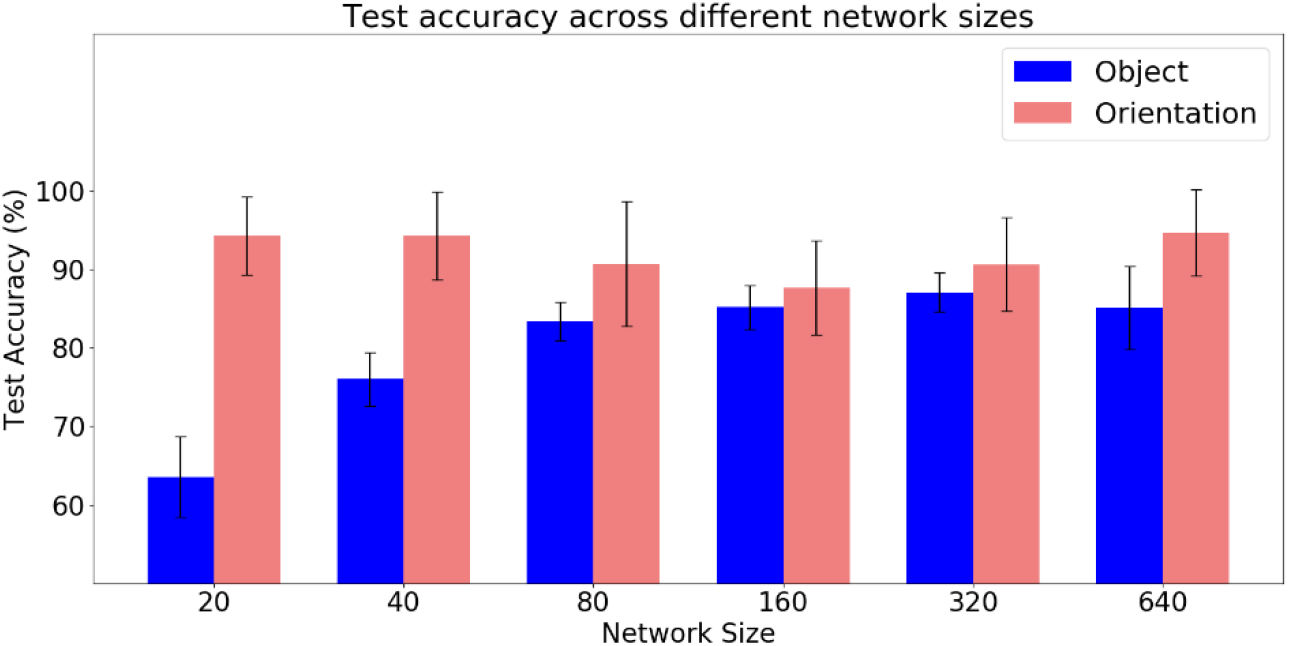
There is a direct relationship between object task’s accuracy and network size in LSTMs trained on the COIL-100. Blue and red bars represent object and orientation classification accuracies, respectively. Error bars are standard errors of the mean (SEM).

**Supplementary figure 2:**
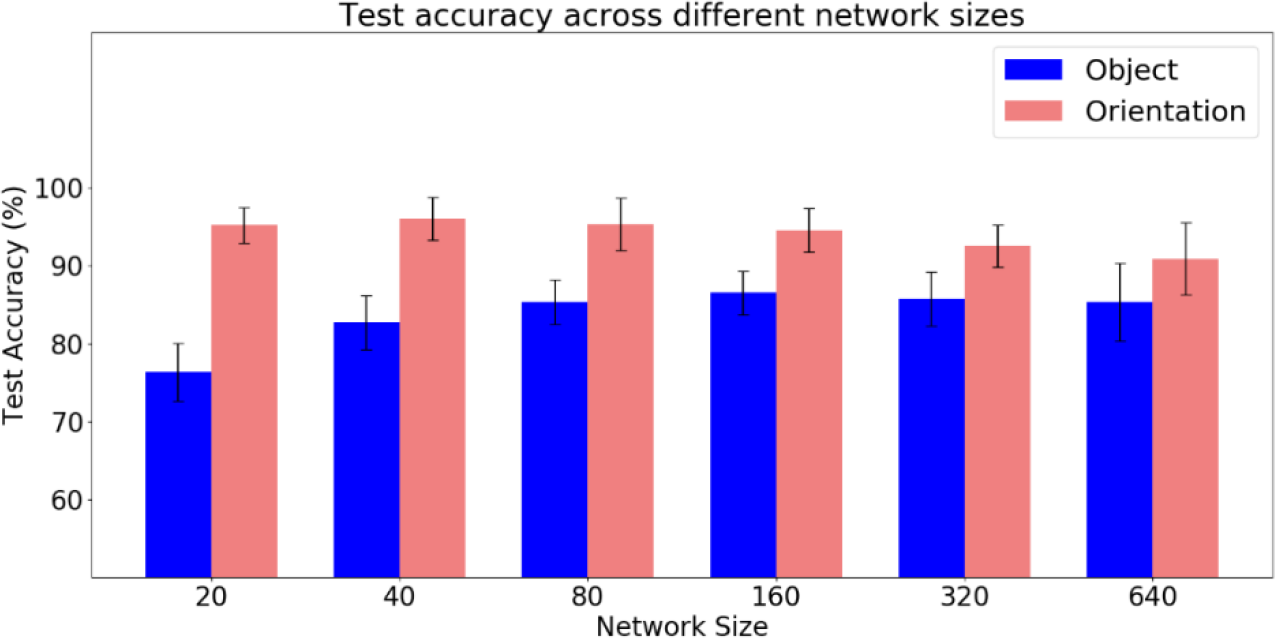
There is a direct relationship between object task’s accuracy and network size in LSTMs trained on the Visual MIRO pairs of tasks. Blue and light red bars represent object and orientation classification accuracies, respectively. Error bars are standard errors of the mean (SEM).

**Supplementary figure 3:**
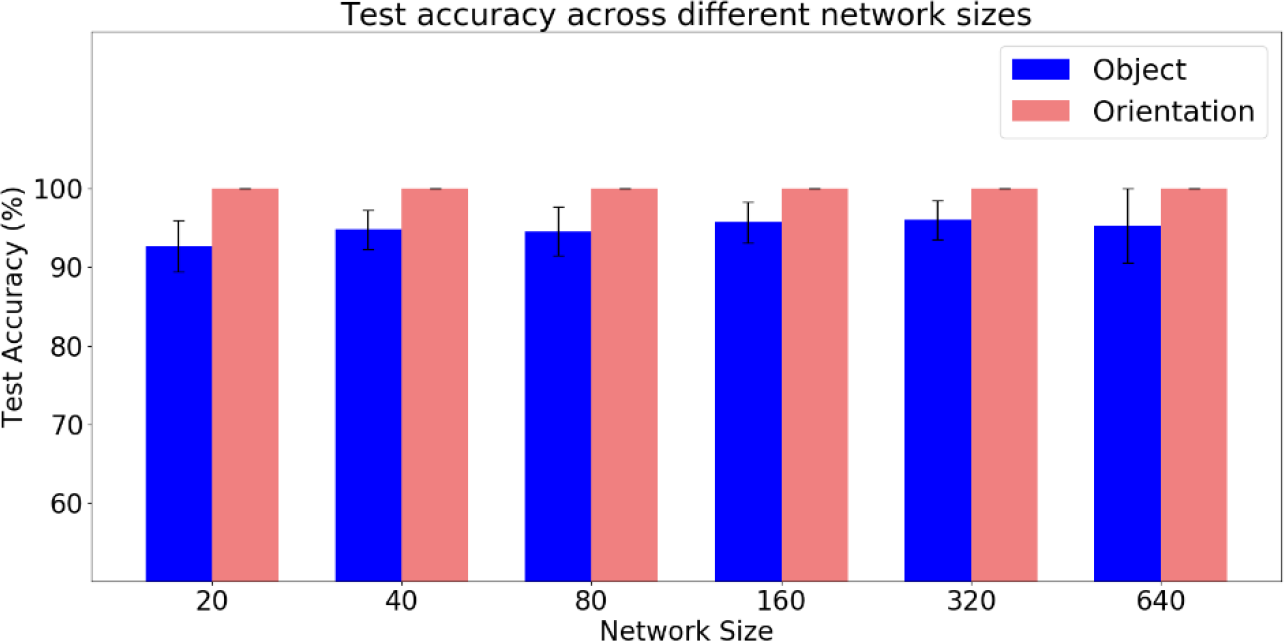
There is a slightly positive relationship between object task’s accuracy and network size in LSTMs trained on the Semantic MIRO pairs of tasks. Blue and light red bars represent object and orientation classification accuracies, respectively. Error bars are standard errors of the mean (SEM).

**Supplementary figure 4:**
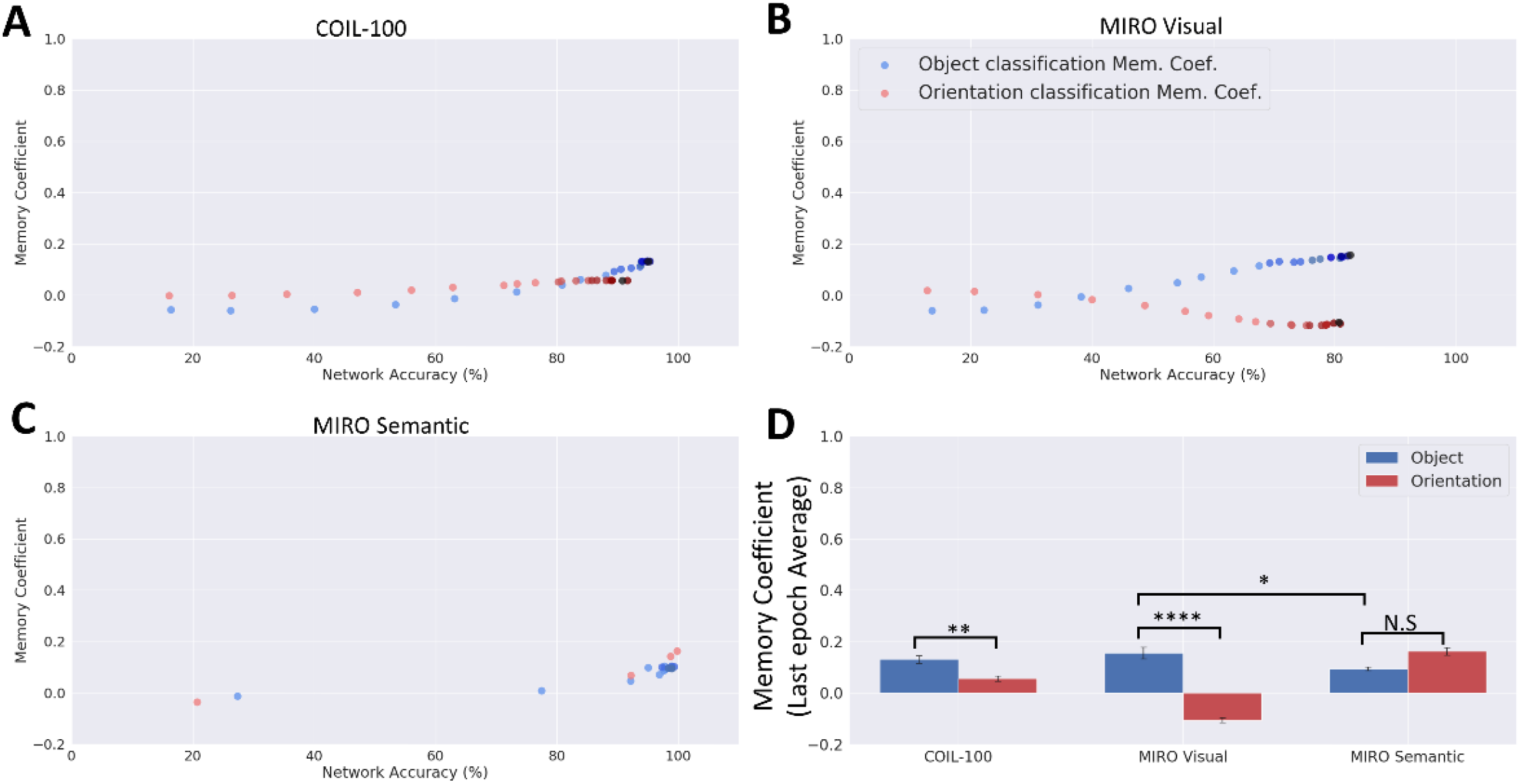
Object task requires a longer memory across COIL-100 and Visual MIRO pair of tasks even when number of classes are equal in both object and orientation tasks. In this experiment, object task had 6 distinct classes (36 images each) that were randomly chosen from the main 54 classes (216 in total). The orientation task had 6 classes constructed from the 216 pool of images used in the object task each containing 36 images of different objects with the same orientation. Orientation angles for defining classes were the same as the main COIL-100 task pairs explained in methods. A) Memory coefficient as a function of task accuracy across learning epochs for the COIL-100 dataset trained on a network size of 20 neurons. Each dot represents the average memory coefficient and accuracy of 30 model realizations. Since there were 200 epochs in LSTM training, only every 10 epochs are shown here (i.e., 1, 11, 21, …, 191, and 200), and darker shades indicate later epochs. Object task requires a longer memory. B) Memory coefficient as a function of task accuracy across learning epochs for the MIRO Visual pair of tasks. Object task still require a longer memory. Object task requires a longer memory. All conventions are the same as A. The object task was defined as 8 object classes (64 images each) that were randomly chosen from the main 22 classes (512 in total). The orientation task had 8 classes constructed from the 512 pool of images used in the object task. Each orientation class contained 64 images of different objects with the same orientation. Orientation angles for defining classes were the same as the main Visual MIRO task pairs explained in methods. C) Same as B but for the MIRO Semantic pair of tasks. Memory dependence is lower when objects are visually dissimilar. Note that the orientation task’s final epochs were epoch 41 where our model reached > 99% test accuracy. The object task was defined as 4 object classes (320 images each) that were randomly chosen from the main 6 classes (1280 in total). The orientation task had 4 classes constructed from the 1280 pool of images used in the object task each containing 320 images of different objects with the same orientation. Orientation angles for defining classes were the same as the main Visual MIRO task pairs explained in methods. D) Average memory coefficient of last epochs over all 30 realizations are shown. Two tasks showed significant differences between object and orientation tasks (*p* < 10^-3^, *p* < 10^-15^, and *p* = .020 for COIL-100, MIRO Visual, and MIRO Semantic tasks, respectively). Moreover, there was a significant difference between object tasks in Visual MIRO pair and the object task in Semantic MIRO pair, suggesting that visual similarity contributes to longer memory (*p* = .010). Error bars show the standard error of the mean (SEM).

**Supplementary figure 5:**
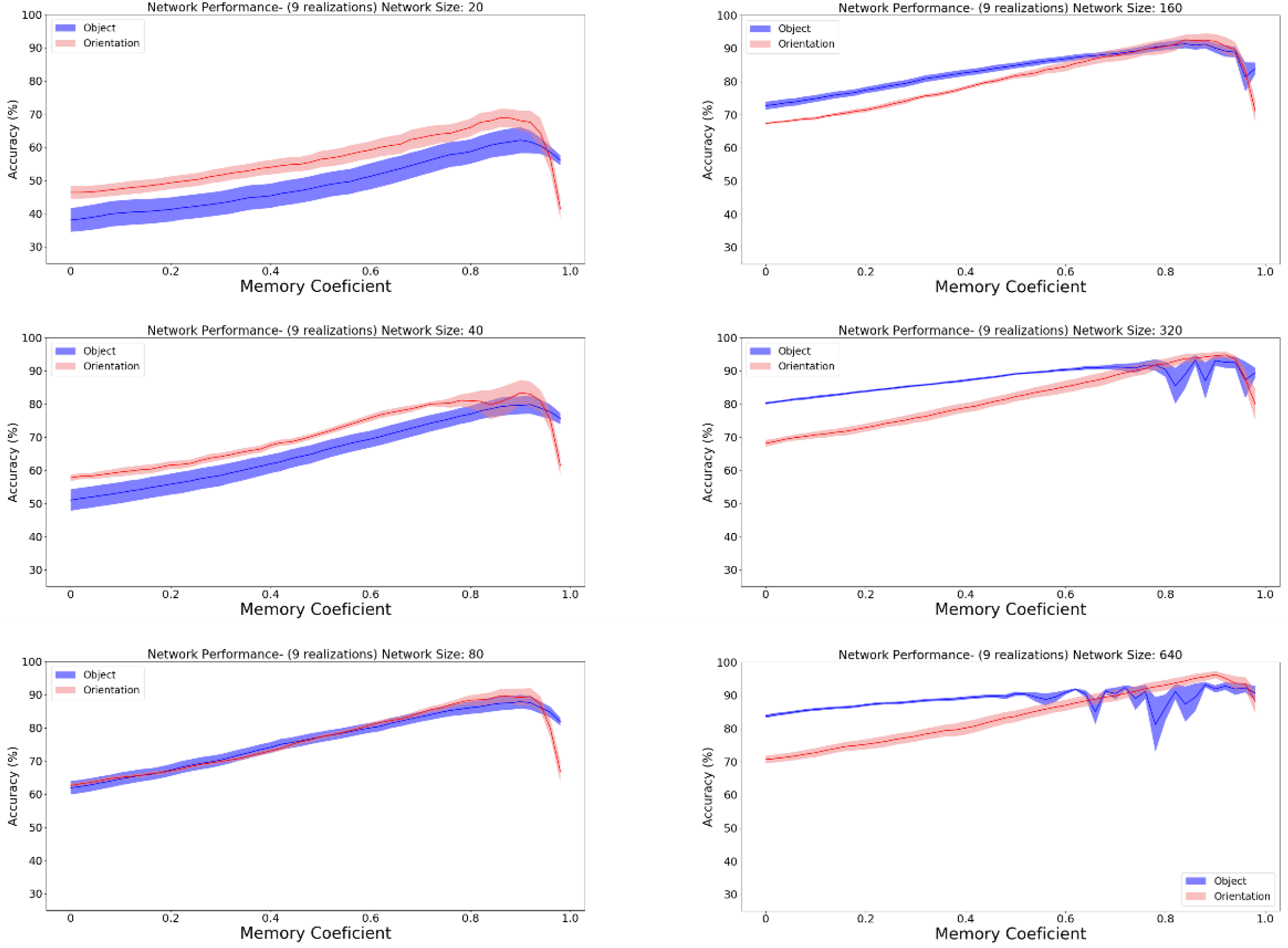
In COIL-100 dataset, memory-accuracy profiles of the two tasks were most similar when the network size was 160. Each subpanel indicates the accuracy of both object (blue) and orientation (red) tasks for a given memory coefficient for a specific network size. Network sizes range from 20 to 640 and memory coefficients range from 0 to 1 with 0.02 spacing. To obtain each accuracy value, the memory coefficients were kept at a constant value and networks were trained (output weights were allowed to adjust by gradient decent) for 50 epochs. After 50 epochs, networks reached a relative performance plateau, and their accuracy was recorded. Results are the average accuracy of 9 different network realizations, and shaded areas are standard error of the mean (SEM). The network size of 160 was chosen to investigate the optimal memory coefficient for each task since at this network size, output weight adjustment affects both tasks similarly and memory coefficient could be studied in isolation.

**Supplementary figure 6:**
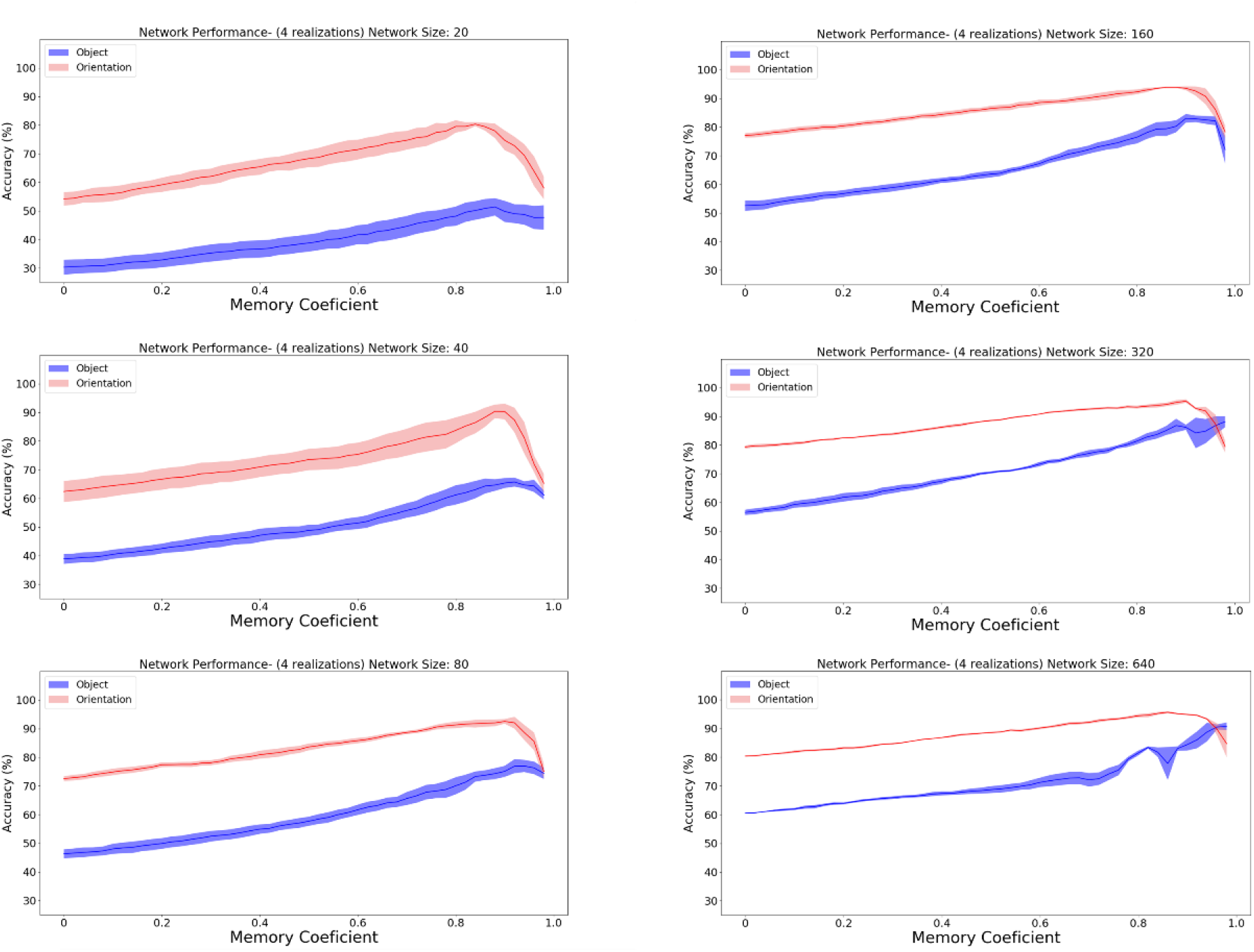
In Visual MIRO pair of tasks, memory-accuracy profiles of the two tasks were most similar when the network size was 160. Each subpanel indicates the accuracy of both object (blue) and orientation (red) tasks for a given memory coefficient for a specific network size. Network sizes range from 20 to 640 and memory coefficients range from 0 to 1 with 0.02 spacing. To obtain each accuracy value, the memory coefficients were kept at a constant value and networks were trained (output weights were allowed to adjust by gradient decent) for 50 epochs. After 50 epochs, networks reached a relative performance plateau, and their accuracy was recorded. Results are the average accuracy of 4 different network realizations, and shaded areas are standard error of the mean (SEM). The network size of 160 was chosen to investigate the optimal memory coefficient for each task since at this network size, output weight adjustment affects both tasks similarly and memory coefficient could be studied in isolation.

**Supplementary figure 7:**
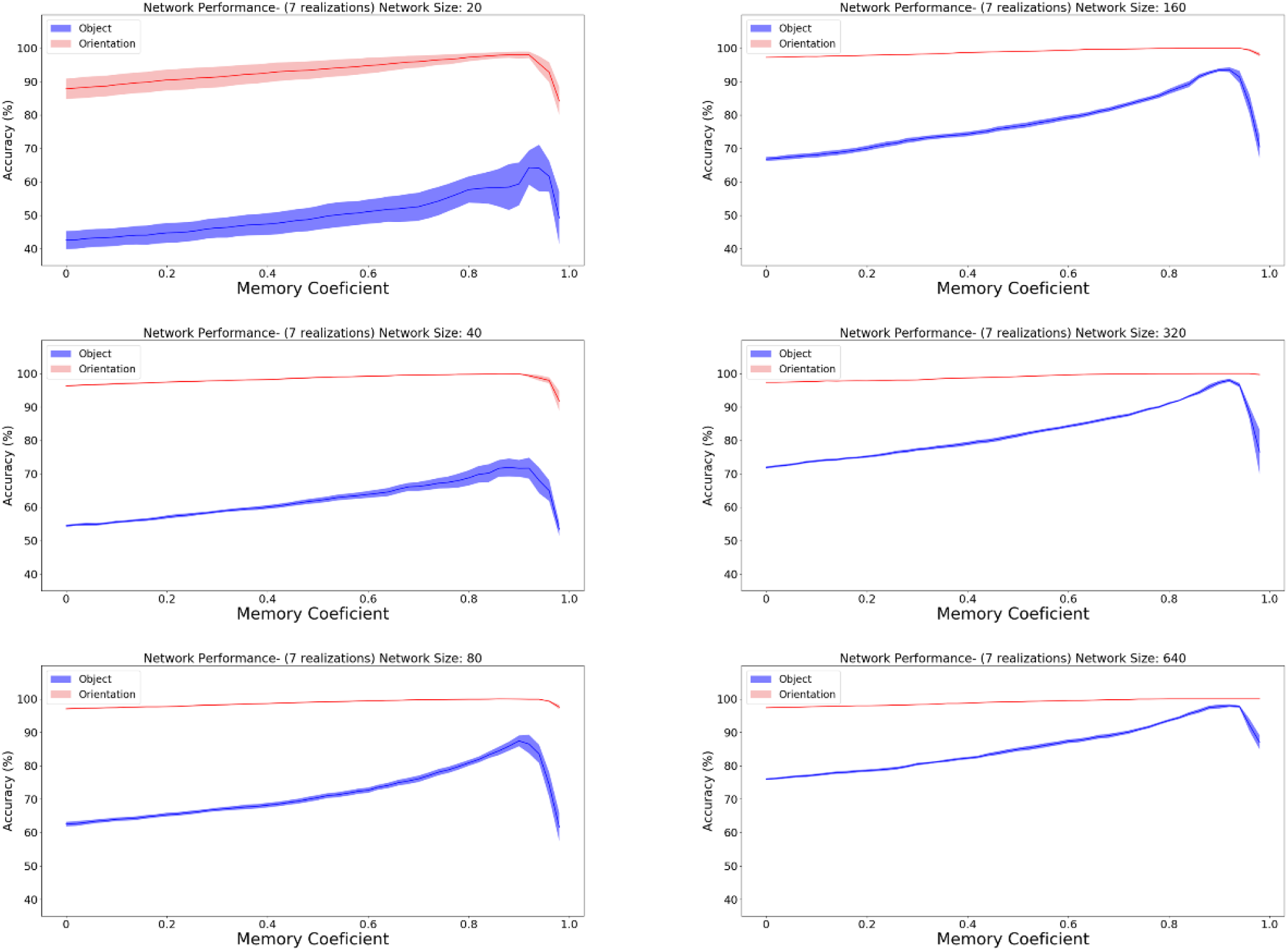
In Semantic MIRO pair of tasks, memory-accuracy profiles of the two tasks were most similar when the network size was 20. Each subpanel indicates the accuracy of both object (blue) and orientation (red) tasks for a given memory coefficient for a specific network size. Network sizes range from 20 to 640 and memory coefficients range from 0 to 1 with 0.02 spacing. To obtain each accuracy value, the memory coefficients were kept at a constant value and networks were trained (output weights were allowed to adjust by gradient decent) for 50 epochs. After 50 epochs, networks reached a relative performance plateau, and their accuracy was recorded. Results are the average accuracy of 4 different network realizations, and shaded areas are standard error of the mean (SEM). The network size of 120 was chosen to investigate the optimal memory coefficient for each task since at this network size, output weight adjustment affects both tasks similarly and memory coefficient could be studied in isolation.

